# Immune Profiling the Axilla with Fine Needle Aspiration is Feasible to Risk-Stratify Breast Cancer

**DOI:** 10.1101/2025.10.22.683956

**Authors:** Jasmine Gore, Amy Llewellyn, Chuen Lam, Jacqueline Shields, Kalnisha Naidoo

## Abstract

Whether breast cancer (BC) has metastasised to the regional axillary lymph nodes (ALN) is an important prognostic factor that directs the extent of axillary surgery. However, surgically clearing the axilla carries morbidity, so less invasive methods of risk-stratifying patients are needed. ALN fine needle aspiration (FNA) is currently used to detect BC metastases, but these samples also contain immune cells. Here we show that FNA acquires sufficient leukocytes for comprehensive immunophenotyping of reactive, patient-derived ALN. All CD4^+^ and CD8^+^ T cells subsets (naïve, terminal effector, central and effector memory) and rarer (<2%) natural killer (NK) and plasmacytoid dendritic cell (pDCs) populations are represented. Importantly, the immune cell profile of one reactive ALN appears to reflect the immune status of the patient’s axilla.

We show that FNA captures immune differences between patients with ≤1 or ≥2 metastatic ALN. Significantly increased numbers of naïve CD4^+^ T cells, but significantly fewer terminal effector, central memory and effector memory subpopulations, were obtained from patients with ≥2 metastatic ALN. Moreover, despite their sparse distribution pattern on whole-section immunohistochemistry (WSI), FNA revealed that CD56^+^ NK cell activation receptors were decreased in patients with ≥2 metastatic ALN. Finally, FNA captured a significant decrease in pDCs in patients with ≥2 metastatic ALN, despite their clustered distribution pattern on WSI.

In summary, FNA is not only feasible for sampling leukocytes from reactive, patient-derived ALN, but also identifies immune cell profiles that reflect axillary tumour burden in BC. Thus, this technique could be used to risk-stratify BC patients in future.

## INTRODUCTION

A third of breast cancer (BC) patients present with axillary lymph node (ALN) metastases [1], and these patients have poorer outcomes [2]. If adjuvant systemic therapy is not given, patients with isolated tumour cells (<0.2 mm or <200 cells) or micrometastasis (0.2-2.0 mm) in ALN have a lower 5-year survival than patients with negative (tumour-free) ALN [3]. Similarly, BC patients with ≥3 metastatic ALN have a worse survival than those with one or two [4, 5].

Sentinel lymph nodes (SLN), which identify the region of the axilla where breast fluid drains, are the first ALN to which BC spreads [6, 7]. Accurately assessing SLN status is essential to treatment planning. ALN that look suspicious on pre-operative imaging are sampled with fine needle aspiration (FNA) or core biopsy to detect metastases [8]. If these samples are negative, a SLN biopsy (SLNB) is performed for histopathological examination during breast surgery [8, 9].

This conservative surgical approach is justified by clinical trials that proved that an ALN clearance (ALNC) is not always necessary. The NASBP B-32 trial showed that in patients with negative SLN, overall and disease-free survival after SLNB alone was equivalent to an ALNC [10]. The Z0011 trial showed the same for patients with only one or two micrometastatic SLN [5]. Clinically however, deciding whether patients with one or two macrometastatic (>2mm) ALN require a completion clearance remains contentious [11]. Recently, the INSEMA and SOUND trials have challenged if ALN surgery is necessary at all in early-stage BC (T1 and/or T2 with a clinically negative axilla) by showing that omitting ALN surgery is non-inferior to SLNB when assessing disease-free survival [12, 13]. Additionally, the emergence of checkpoint inhibitors together with the understanding that immune system regulates tumour progression has prompted further research into how/when it is safe to omit a SLNB in those patients who are not at risk of axillary spread.

The LN is an immunological hub that surveys for pathogens and immune threats, and its composition reflects the pathophysiology of the tissues it drains [14]. In pre-metastatic LN, alterations in B- and T-cell localisation and relative abundance of T-cell subsets is thought to precede the arrival of cancer cells [15]. ALN immunophenotype also has been shown to change in advanced BC patients [16], and in murine metastatic nodes, T-cells can move towards an effector/memory and exhausted phenotype [17]. Thus, it may be beneficial to look beyond the tumour and its metastases, into the LN response itself, to identify immunological signatures that predict disease progression.

While FNA is used diagnostically to detect metastatic tumour cells in ALN, these samples also contain immune cells. Using FNA with downstream single-cell RNA-sequencing, Provine *et al.* detected T-cells, B-cells, natural killer (NK) cells, monocytes, plasmacytoid dendritic cells (pDC) and conventional dendritic cells in cervical LN [18]. Similarly, in non-small cell lung carcinoma patients, FNA sampling identified site-specific T-cell changes, showing significantly fewer CD4^+^, and more regulatory, T-cells in tumour-draining versus non-tumour draining nodes [19]. Finally, immunoprofiling of joint-draining inguinal LN FNA samples revealed significant differences in B- and CD4^+^/CD8^+^ T-cell composition between healthy volunteers, patients with early rheumatoid arthritis (RA) and individuals at risk of developing RA [20–22]. This suggests that FNA immunoprofiling can be used in early disease to predict outcome/risk, but this has yet to be tested in BC.

Here, we sought to determine if FNA can be used to comprehensively immunoprofile BC patient-derived ALN and stratify patients according to axillary tumour burden. We confirm that FNA are rich in, and can detect, diverse immune populations in reactive, patient-derived ALN. Furthermore, preliminary data shows that FNA is sensitive enough to detect distinct immune changes between patients with one or two metastatic ALN.

## RESULTS

### FNA samples the same immune cells as whole LN digestion

To first determine if FNA captured the immune diversity seen in standard whole node processing methods, we compared the contents of FNA samples to whole LN cell suspensions (i.e. mechanical disruption or enzymatic digestion with either Type 1 collagenase or collagenases A&D) using murine LN. The number and viability of immune cells in FNA samples was comparable to those obtained with collagenase digestion (Fig. 1A and B, fig. S1); mechanical digestion was the least efficient (*n = 2* (zero cells collected from one LN)).

**Fig. 1.**
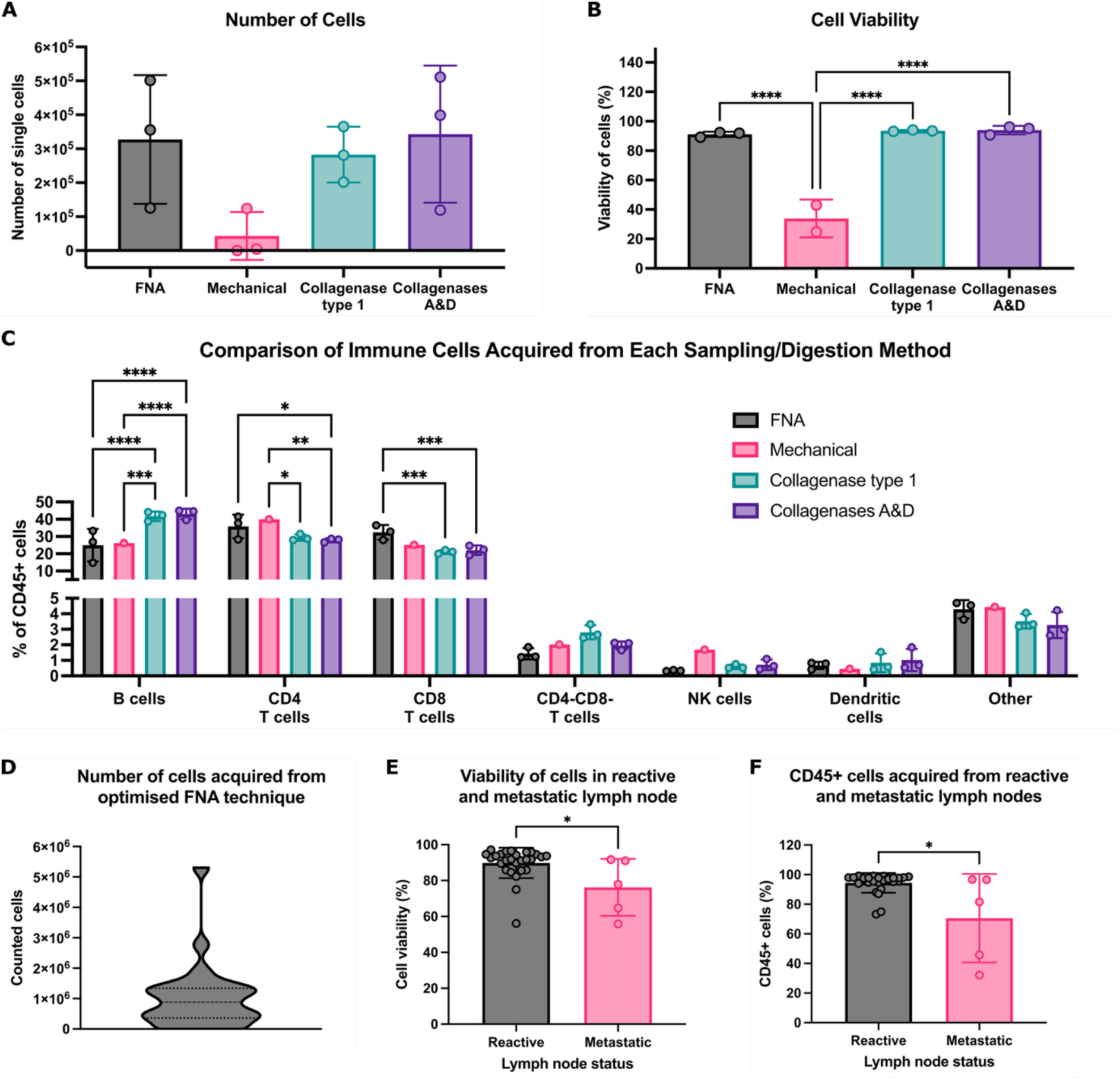
FNA reproducibly captures diverse immune cell populations in mice and is feasible in patient-derived samples. (**A**) Comparison of the number of cells acquired between LN sampling methods in C57/BL6 female mice, quantified by flow cytometry. Data presented as mean ± SD, each point represents one inguinal LN*, n = 3* per group. One-way ANOVA was used to determine statistical significance. (**B**) Quantification of cell viability across LN sampling methods in mice. Data presented as mean ± SD, each point represents percentage viability of cells from one inguinal LN, *n = 3* per group apart from mechanical digestion where *n = 2*. One-way ANOVA was used to determine statistical significance. (**C**) Quantification of immune cell abundance obtained between each sampling method in mice. Data presented as mean ± SD, each point is one inguinal LN, *n = 3* per group apart from mechanical digestion where *n = 1*. Two-way ANOVA used to determine statistical significance. (**D**) Distribution of the number of cells acquired from the optimised FNA technique in patient samples. Cells were counted manually on a haemocytometer immediately post sampling; *n = 27* ALN. (**E**) Comparison of cell viability between reactive and metastatic patient-derived ALN measured by flow cytometry. Data presented as mean ± SD. Reactive ALN; *n = 27*, Metastatic ALN; *n = 5*. Mann-Whitney Test was used to determine significance. (**F**) Comparison of the percentage of CD45^+^ cells acquired between reactive and metastatic patient-derived ALN determined by flow cytometry. Data presented as mean ± SD. Reactive ALN; *n = 27*, Metastatic ALN; *n = 5*. Mann-Whitney Test was used to determine significances. (\**P* <.05, \*\**P* <.01, \*\*\**P* <.001, and \*\*\*\**P* <.0001).

Importantly, FNA sampled all the immune cell (CD45^+^) populations within the node (Fig. 1C). B-cells were abundant in FNA, albeit at lower levels than collagenase digestion methods. Conversely, FNA enriched for CD4^+^/CD8^+^ T-cell populations. Importantly, rarer populations (CD4^-^CD8^-^ T-cells, NK cells and/or DC) were detected in similar proportions by both digestion and FNA. Interestingly, while enzymatic digestion identified positive and negative cell populations, these were dimmer and less distinct than FNA. indicating that FNA sampling negated the effect of enzymatic digestion on cell surface epitopes (fig. S2).

Thus, FNA can acquire abundant, diverse immune populations from murine LN without impacting cell viability or epitope availability.

### FNA is feasible and identifies diverse immune populations in patient-derived ALN

We then tested feasibility in patient ALN. On average, FNA acquired 1 x 10^6^ cells per reactive ALN (Fig. 1D). Interestingly, the presence of ALN metastasis significantly decreased cell viability (Fig. 1E); *P* = .02) and significantly fewer immune cells (CD45^+^) were acquired from metastatic ALN (Fig. 1F; (*P* = .03)).

With FNA, we could identify and quantify the major immune cell populations within reactive ALN (Fig. 2A and B, and fig. S3A; gating strategy shown in fig. S4; *n = 9* patients). As expected, most were T-, and B-, cells. We could also detect and characterise rarer populations e.g. pDC and NK cells. Moreover, FNA discriminated between immune cell subpopulations within each cluster (Fig. 2C–E) detecting naïve, terminal effector, central memory and effector memory T-cell subsets, alongside identifiers of functional and activation state. Finally, we compared immune cell proportions between patients and demonstrated limited inter-nodal variability (fig. S3B).

**Fig. 2.**
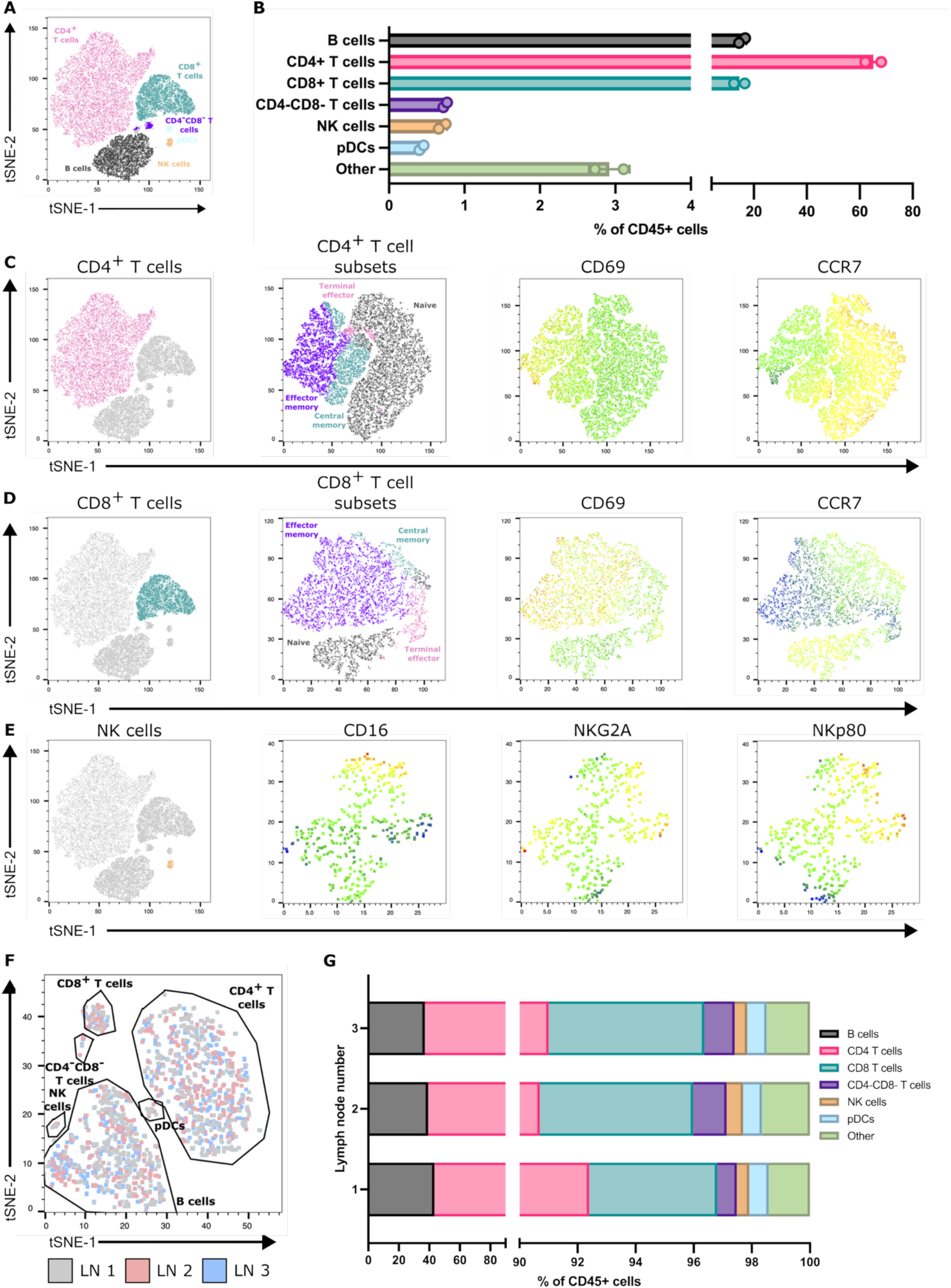
FNA consistently acquires diverse immune populations across reactive, patient-derived ALN. (**A**) Representative flow cytometry derived t-SNE on CD45^+^ cells from one patient sample showing the major immune cell populations acquired. (**B**) Quantification of immune cell populations detected within FNA samples from the same patient. Data presented as mean ± SD. Each point represents one ALN; *n =2.* (**C**) Representative t-SNE showing CD4^+^ T-cell subsets detected within samples; naïve, black; terminal effector, pink; effector memory, purple; central memory, aqua) and CD69 and CCR7 expression within these subsets (colour scale - blue, low expression; red high expression), (**D**) Representative t-SNE showing CD8^+^ T-cell subsets and CD69 and CCR7 expression, and (**E**) NK cell surface receptor expression identified through FNA sampling. (**F**) Representative t-SNE generated from flow cytometry data depicting overlapping immune cell clustering from each of three ALN collected from the same patient. (**G**) Flow cytometry quantification of proportions of immune cell populations between each of the three ALN from the same patient.

### One reactive ALN represents the immune status of the axilla

Typically, one to four ALN are collected during a SLNB, and more during an ALNC [23]. Therefore, we assessed the degree of variability in immune cell composition between individual, reactive ALN from each patient. T-SNE analysis showed that all reactive ALN from the same patient clustered consistently, irrespective of where the node is in the chain (Fig. 2F and G; fig. S5). Thus, all reactive ALN within a patient have a similar immune profile.

### FNA and immunohistochemistry of the same ALN show similar immune profiles

Immunohistochemistry (IHC) is used routinely in diagnostic practice to immunophenotype patient-derived ALN. Therefore, we compared FNA sampling with IHC of an abundant cell population (CD4^+^ T-cells), as well as rarer cell populations (CD56^+^ NK cells or CD123^+^ pDC), from the same ALN to ensure that FNA sampling was representative (*n = 8* patients).

IHC quantification and FNA identified similar proportions of CD4^+^ T-cells (Fig. 3A and B).

**Fig. 3.**
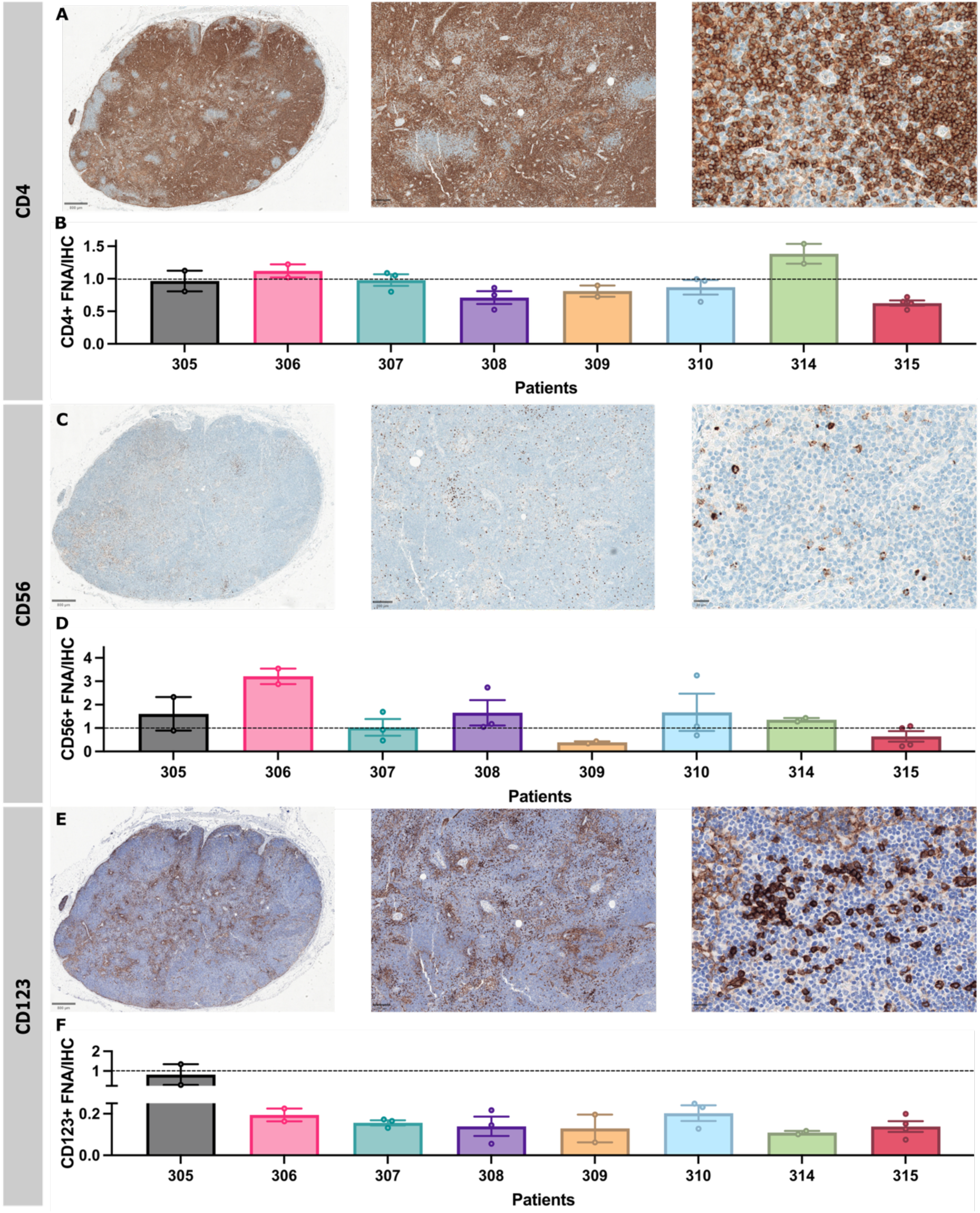
FNA and IHC of the same ALN identify similar abundant and scarce immune cell types within patient-derived ALN. (**A**) Representative photomicrographs of patient-derived ALN immunostained with CD4 (birds-eye view, x50 (inset i) and x400 (inset ii) magnification. (**B**) Proportion of CD4^+^ cells in FNA sample compared to that in IHC images. (**C**) Representative photomicrographs of patient-derived ALN immunostained with CD56 (birds-eye view, x50 (inset i) and x400 (inset ii) magnification. (**D**) Proportion of CD56^+^ cells in FNA sample compared to that in IHC images. (**E**) Representative photomicrographs of patient-derived ALN immunostained with CD123 (birds-eye view, x50 (inset i) and x400 magnification (inset ii). Samples were counterstained with Haematoxylin for nuclear identification. (**F**) Proportion of pDCs (CD4^+^HLA^-^DR^+^) in FNA sample compared to proportion of that in IHC images. All bars are mean ± SD. Each point represents one ALN.

On IHC, NK cells showed strong, granular cytoplasmic CD56 staining and were singly dispersed throughout the paracortex (Fig. 3C). There was some variation in the FNA proportions of NK cells compared to IHC-based image analysis from individual patients - FNA generally enriched for NK cells, but only patient 306 was a significant outlier (Fig. 3D). pDC showed strong cytoplasmic and membranous CD123 staining and were patchily distributed as clusters throughout the paracortex (Fig. 3E). High endothelial venules (HEV) and sinus macrophages were weakly CD123 positive. These were excluded during quantification by thresholding for staining intensity. FNA samples contained fewer pDC than IHC (Fig. 3F). Overall, FNA still captured these rarer cell populations, and detected variations in cell surface protein expression in a representative and interpretable fashion.

### Immune cell proportions reflect axillary tumour burden

Finally, we tested if FNA sampling could identify immune differences in reactive ALN from patients with different tumour volumes within their axillae. Patients were stratified into two groups (≤1 or ≥2 metastatic ALN). There were no proportional differences in CD4^+^ and CD8^+^ T-cells or NK cells, but pDC were significantly decreased in the ≥2 metastatic ALN cohort (Fig. 4A; *P =* .03; gating strategy shown in fig. S6).

**Fig. 4.**
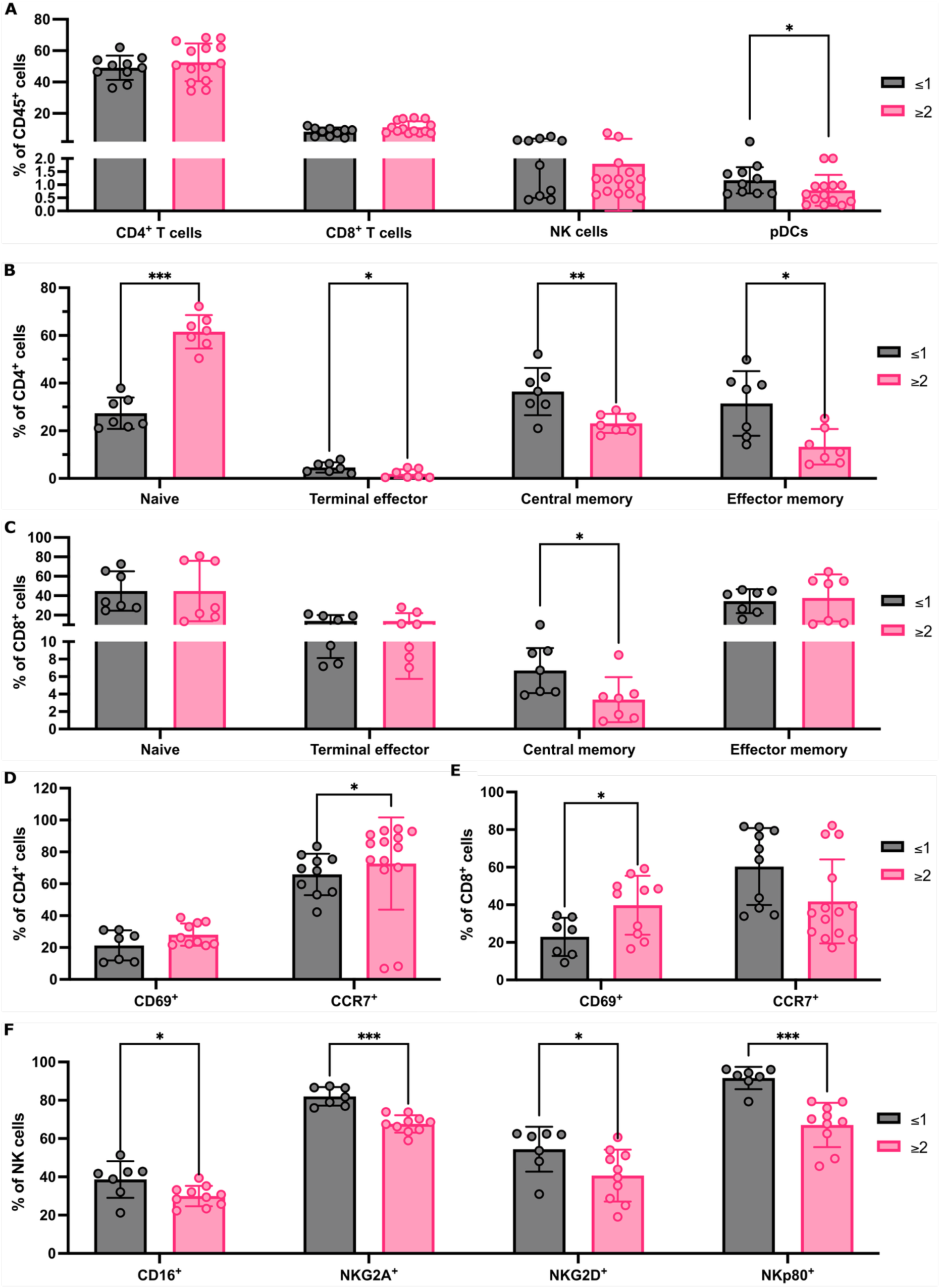
FNA captures immune cell population changes as tumour burden increases in the axilla. (**A**) Flow cytometry quantification of major immune population abundance within FNA samples from patients with ≤1 or ≥2 metastatic ALN. (**B**) Flow cytometry quantification of changes to CD4^+^ T-cell subpopulation abundance between patients with ≤1 or ≥2 metastatic ALN; Naive, terminal effector, central memory and Effector memory subsets presented as % of total CD4 T cells (**C**) Quantification of CD8^+^ T-cell subsets as for (b) between patients with ≤1 or ≥2 metastatic ALN. (**D-E**) Flow cytometry quantification of markers of activation status, CD69 and CCR7, in CD4^+^ **(D)** and CD8^+^ **(E)** T cells comparing patients with ≤1 or ≥2 metastatic ALN. Data represented as % of parent population (CD4 or CD8 respectively). (**F**) Quantification of the effect of total number of metastatic ALN on the frequency of NK cells expressing CD16, NKG2D, NKG2A and NKp80 markers of activation status. Data presented as mean ± SD. £2 ALN; *n ³7*, ³1 ALN; *n ³7,* and each point represents an individual ALN. Mann-Whitney test was used to determine statistical significance. (\**P* <.05, \*\**P* <.0 and \*\*\**P* <.001).

CD4^+^ and CD8^+^ T-cell subpopulations differed between the two groups (Fig. 4B and 4C, Naïve CD4^+^ T-cells (*P* < .001) increased significantly, and CD4^+^ terminal effector (*P* = .04), CD4^+^ effector memory (*P* = .01) and CD4^+^ central memory T-cells (*P* = .007) significantly decreased, in the ≥2 metastatic ALN group. Within CD8^+^ T-cell subsets, central memory T-cells decreased significantly in the ≥2 metastatic ALN cohort (*P* = .01).

We then compared activation receptor CD69 and chemokine receptor CCR7 expression in CD4^+^ and CD8^+^ T-cells (Fig. 4D and 4E). A significant increase in CD69-expressing CD8^+^ T-cells (*P* = .04) and CCR7-expressing CD4^+^ T-cells (*P* = .04) was detected in patients with ≥2 involved ALN, which correlates with the increase in naïve CD4^+^ T-cells observed.

Finally, we evaluated if there were any changes in NK cell phenotype and activation state between groups (Fig. 4F). A significant decrease in CD16 (*P* =.05), NKG2A (*P* < .001), NKG2D (*P* = .03) and NKp80 (*P* < .001) expression was seen in patients with ≥2 metastatic ALN. Together, these data indicate that FNA is sensitive enough to detect distinct immune changes between patients, based on the extent of ALN metastatic disease.

## DISCUSSION

Although SLNB alone is now standard of care for clinically node-negative BC patients, we have yet to define exactly when in early BC the tumour-draining ALN become immunosuppressed, facilitating metastasis. To this end, we have shown that the minimally invasive FNA technique can be used to comprehensively immunophenotype ALN and potentially risk-stratify patients.

The murine experiments confirmed that FNA sampling was representative of whole LN composition. Furthermore, enzymatic digestion commonly alters cell surface receptors and mechanical digestion often fails to isolate DC [24, 25]. FNA avoided both these pitfalls.

Consistent with previous clinical studies using ultrasound-guided FNA, we acquired on average 1 x 10^6^ cells per FNA from patients [26, 27]. Furthermore, our yields were sufficient to comprehensively immunophenotype ALN. The FNA samples from reactive, patient-derived ALN contained a wide range of immune cells, including less abundant populations. As in haematological malignancies [28], FNA acquired more T-than B-cells. However, both B- and T-cell subpopulations were detectable, and functional markers reflected activation state. This carried forward to rarer pDC and NK populations. Even though IHC showed more pDC than matched FNA samples, the latter could still be quantified. Importantly, our data suggests that one reactive ALN is representative of the entire axilla. Translationally, this is relevant as the radiologist could sample the most accessible reactive ALN if, moving forward, FNA were used to risk-stratify patients.

Using IHC, López *et al*. showed that CD8^+^ T-cells and CD68^+^ macrophages significantly increased, but CD123^+^ pDC significantly decreased, in the reactive ALN of node-positive, treatment-naïve BC patients [29]. Interestingly, we have also shown that pDC significantly decrease in the reactive ALN of BC patients once ≥2 ALN elsewhere in the axillary chain contain metastatic tumour. This suggested correlation in the proportion of pDC to axillary tumour burden warrants further investigation in a larger cohort of patients - we plan to do this in future.

We have also shown that NK cell surface expression of CD16, NKG2A, NKG2D and NKp80 correlates with the degree of ALN involvement. Using LN fragments, Frazao *et al*. showed that reactive and metastatic ALN expressed similar NK receptors, but both NKG2A and CD16 expression was significantly increased on NK cells from tumour-draining ALN compared to mesenteric nodes from healthy donors, irrespective of whether they contained tumour, in stage IIIA compared to stage II patients [30]. One could argue that LN from different body sites (in this case, axilla versus abdomen) will always have different immune profiles since they are exposed to different antigens, and therefore mesenteric LN may not have been a suitable control [14, 31]. Moreover, while that study showed that CD16 and NKG2A expression increased in N2-3 versus N1 patients, it did not stratify the N1 cohort based on the number of metastatic ALN as we have.

T-cell researchers have also only compared reactive to metastatic ALN. Vahidi *et al*. showed an increase in central memory CD8^+^, but a decrease in naïve CD4^+^, T-cells in metastatic compared to reactive ALN fragments [32]. This differs from our FNA-based data which showed a decrease in central memory T-cells (including central memory CD4^+^ T-cells), but an increase in naïve CD4^+^ T-cells, in patients with ≥2 involved nodes. This again suggests that the presence of tumour in other nodes within the axillary basin influences the immune response in reactive ALN and are more nuanced than previously thought.

In summary, we have shown that FNA, which is minimally invasive, identifies immune cell changes in reactive, patient-derived ALN in BC. Furthermore, since FNA is already used to diagnose BC nodal metastasis pre-operatively, it could easily be adapted to sample the most accessible, reactive ALN to delineate the patient’s locoregional immune response. Finally, we have shown through a preliminary cohort of BC patients that distinct immunological changes reflect axillary tumour burden; this could be harnessed to risk-stratify patients with early BC in future.

## METHODS

### Patients

ALN tissue samples were collected from BC patients (18–95 years old) undergoing a routine ALNC at King’s College Hospital through the Breast Cancer Immunity, Drug and Gene study (REC reference: 24/NW/0079). Informed consent was obtained from all participants (*n = 10*; Table 1). Samples were de-identified before being released to researchers to ensure blinded data analysis.

**Table 1.** Clinical characteristics for all patients.

| Characteristic | Number of patients (%) |
| --- | --- |
| **Age** |  |
| <50 | 3 (30%) |
| <sup>3</sup>50 | 7 (70%) |
| **Treatment** |  |
| Naïve (Rx) | 6 (60%) |
| NACT | 4 (40%) |
| **Molecular subtype** |  |
| ER<sup>+</sup> | 10 (100%) |
| HER2<sup>+</sup> | 4 (40%) |
| **Number of reactive LN** |  |
| 0 | 1 (10%) |
| <sup>3</sup>1 | 9 (90%) |
| **Number of involved LN** |  |
| £1 | 3 (30%) |
| <sup>3</sup>2 | 7 (70%) |

### FNA of patient-derived ALN

ALN were dissected out of the ALNC specimen by a Histopathologist (AML/KN) at room temperature within one hour of resection. One to six ALN (reactive or metastatic) per patient underwent FNA. While applying a vacuum to a 1mL syringe pre-filled with 500µL phosphate buffered saline (PBS), the attached 23G needle was inserted the ALN for single-pass, multiregional sampling before being withdrawn. The PBS was expelled and the syringe then was reflushed with 500µL of PBS. Cells were counted manually post-sampling using a haemocytometer. Red blood cells within samples were lysed during fixation for flow cytometry. Samples were kept on ice until staining.

### Murine LN Sampling

Charles River female wild-type C57BL/6 mice (*n = 6*) were housed under pathogen-free conditions with a 12-hour day/night cycle, with free access to food and water. Experiments were performed under Project Licence PP2472580. After culling, inguinal LN were removed. FNA were acquired by inserting a 25G needle, attached to a 1 mL syringe pre-filled with 500µL, into excised nodes (*n = 3*). The needle was twisted while simultaneously pulling on the syringe plunger during a single pass. Fluid was then expelled from the syringe. Whole inguinal LN (*n = 3* per condition) were enzymatically digested with Type 1 collagenase or collagenases A&D (both 1 mg/mL), and DNase (0.1 mg/mL) for 30 mins at 37 °C at 500 rpm. Suspensions were pipetted at 15 mins to aid digestion. Reactions were stopped with 20 mM ethylenediaminetetraacetic acid. Cells were passed through 100mm cell strainer which was subsequently washed with PBS. For mechanical disruption, whole inguinal LN (*n = 3*) were teased apart using 25G needles on a 100mm cell strainer. The LN was mashed with the end of a 1mL syringe plunger into a single cell suspension on the strainer; both were then rinsed with 10mL of PBS. Only two LN had enough cells to assess cell viability and only one could be immunophenotyped.

### Flow cytometry

Suspensions were centrifuged at 500g for 5 mins and washed with PBS, then stained with Zombie UV™ Live Dead dye (Biolegend) at 1:500 in PBS for 30 mins in the dark, room temperature. Cells were then washed with FACS buffer (2% foetal bovine serum (FBS) in PBS) and resuspended in blocking antibody at 1:33 (human) or 1:500 (mouse) for 15 mins, room temperature (human) or 4°C (mouse). Chemokine receptor antibodies were added to samples at 1:200, incubated at 37°C for 30 mins before remaining antibodies (tables S1 and S2) were added at 1:200 and incubated for 1 hour, 4°C. Samples were washed and fixed (FoxP3/Transcription Factor Fixation/Permeabilisation, eBioscience™) for 15 mins at room temperature which lysed any red blood cells. Cells were then permeabilised and incubated with blocking antibody in permeabilisation buffer (FoxP3/Transcription Factor Fixation/Permeabilisation, eBioscience™) for 15 mins, room temperature. Intracellular antibodies were added at 1:200 for 30 mins, 4°C. Samples were then washed and resuspended with FACS buffer and read on either CytoFLEX (Beckman Coulter) or LSRFortessa (BD Biosciences). During analysis, FNA that acquired fewer than 1x10^4^ CD45^+^ cells were excluded from immunophenotyping analysis. Data was analysed using FlowJo 10.10.0. A native platform within FlowJo was used to generate t-SNEs.

### IHC

Formalin-fixed and paraffin-embedded reactive ALN tissue sections (3mm thick; *n = 8* patients) were immunostained using automated staining protocols with either CD4 (1:80), CD56 (1:50) or CD123 (1:100) antibodies after heat-induced epitope retrieval in pH9 buffer. Automated staining was completed using the Optiview Kit on Roche Benchmark Ultra for CD4 and CD56, or BOND Compact Polymer Detection kit on Leica Bond III for CD123.

### Image analysis

Whole-slide images were captured using the Glissando Desktop Scanner (Objective Imaging). The proportion of CD4^+^, CD56^+^ and CD123^+^ cells was determined using QuPath software V.0.6.0 by quantification of the whole ALN (fig. S7) [33]. The pixel classifier using a hematoxylin stain threshold was used to exclude background areas. A second pixel classification for DAB staining was used on CD123 images to exclude weakly positive HEV and sinuses, confirmed with a manual adjustment by a Histopathologist (AML/KN). IHC-positive cells were calculated as a percentage of all cell nuclei in the selected ALN areas using the positive cell detection tool. FNA-positive cells were calculated as a percentage of all single cells and compared to IHC-positive cells as a percentage of all cell nuclei (CD4^+^ and CD56^+^) or cell nuclei outside of HEV and sinuses (CD123^+^) in tissue sections.

### Statistical analysis

All statistical analysis was performed on GraphPad Prism V 10.4.1. For parametric data, two groups were compared by One-Way ANOVA; > two groups were analysed by Two-Way ANOVA. Pairs of unmatched, non-parametric data were compared using the Mann-Whitney test. Kruskal-Wallis Test was used for multiple comparisons of non-parametric data. A p-value of <0.05 was considered significant.

## DATA AVAILIBITY

Data in this study can be made available from the corresponding authors on request.

## CODE AVAILABILITY

Code used in this study is publicly available using the QuPath platform.

## ACKNOWLEDGEMENTS

The authors would like to thank the patients for agreeing to donate their tissue samples to this study. The authors would like to thank members of the KCL New Hunts House BSU facility and for technical assistance and support. The authors would also like to thank members of the Breast Cancer Care Team at King’s College Hospital for their help in co-ordinating tissue sample collection.

## CONFLICT OF INTEREST STATEMENT

Authors declare that they have no conflict of interests.

## FUNDING

This work was supported by Cancer Research UK (EDDPJT-Nov22/100036) (JDS).

## ETHICAL APPROVAL

Patients were recruited through the Breast Cancer Immunity, Drug and Gene (BRIDGE) study (clinical trial number: not applicable; approved by the HRA and Health and Care Research Wales (Research Ethics Committee reference: 24/NW/0079)) on 24 March 2024. Informed consent was obtained from all participants by trained members of the research team either in person or over the phone. Study participants could withdraw consent at any time during the study period.

Mouse experiments were performed under Project Licence PP2472580 approved on 6 April 2023. Mice were housed under pathogen-free conditions with a 12-hour day/night cycle, with free access to food and water.

## AUTHOR CONTRIBUTIONS

JS and KN conceived the project. JS, KN, JG designed the methodology. JG performed *in vivo* experiments. JG, AL and KN obtained FNA samples. JG processed samples, stained and analysed flow cytometry data. CL stained tissue sections. AL and KN examined stained tissue sections, which were quantified by JG. JG wrote original draft, which was reviewed and edited by KN and JS. JS, KN and JAG were responsible for project administration. JS acquired funding, KN and JS supervised the research.

## Supplementary Materials

### Supplementary Figures

**Figure S1.**
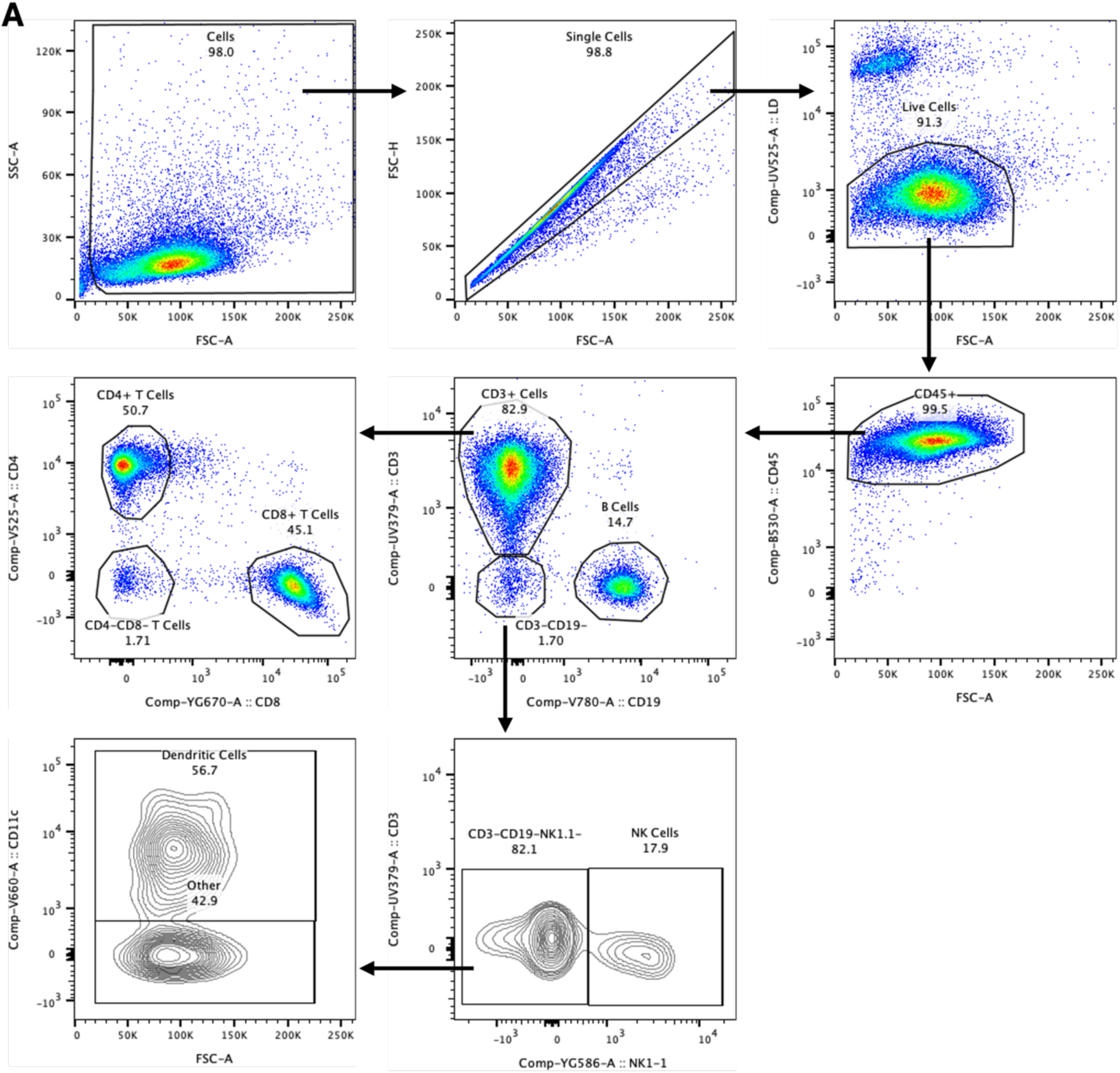
Gating strategy to identify murine immune cell populations in inguinal LN.

**Figure S2.**
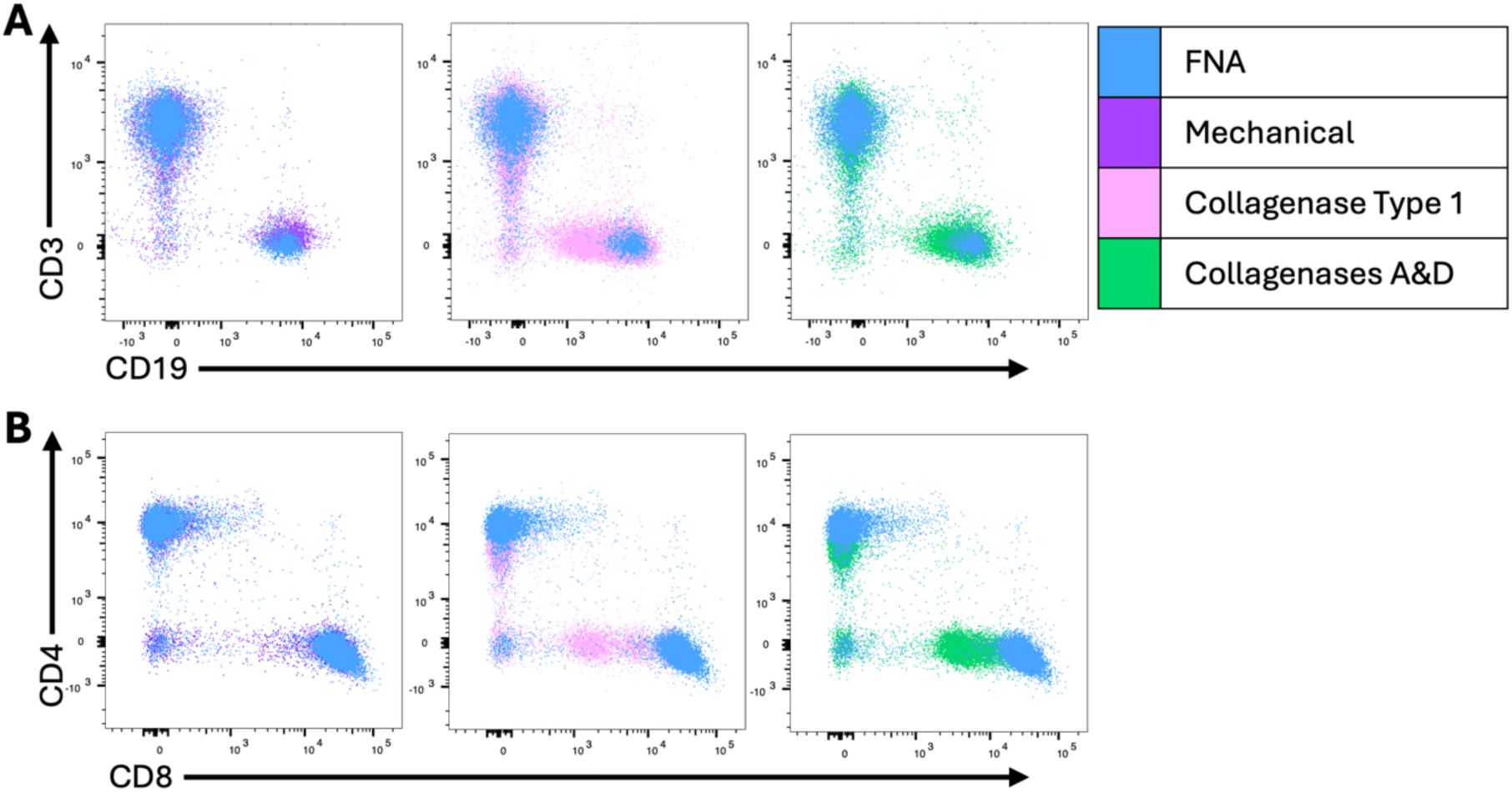
FNA preserves the integrity of cell surface markers CD3, CD19, CD4 and CD8. Murine inguinal LN were sampled/digested using either FNA, mechanical digestion, collagenase type 1 or collagenases A&D. FNA (blue) is overlaid on mechanical (purple), collagenase type 1 (pink) or collagenases A&D (green). (**A**) Dot plots showing staining of CD3 and CD19 on CD45^+^ immune cells. (**B**) Dot plots showing staining of CD4 and CD8 on CD45^+^CD3^+^ immune cells.

**Figure S3.**
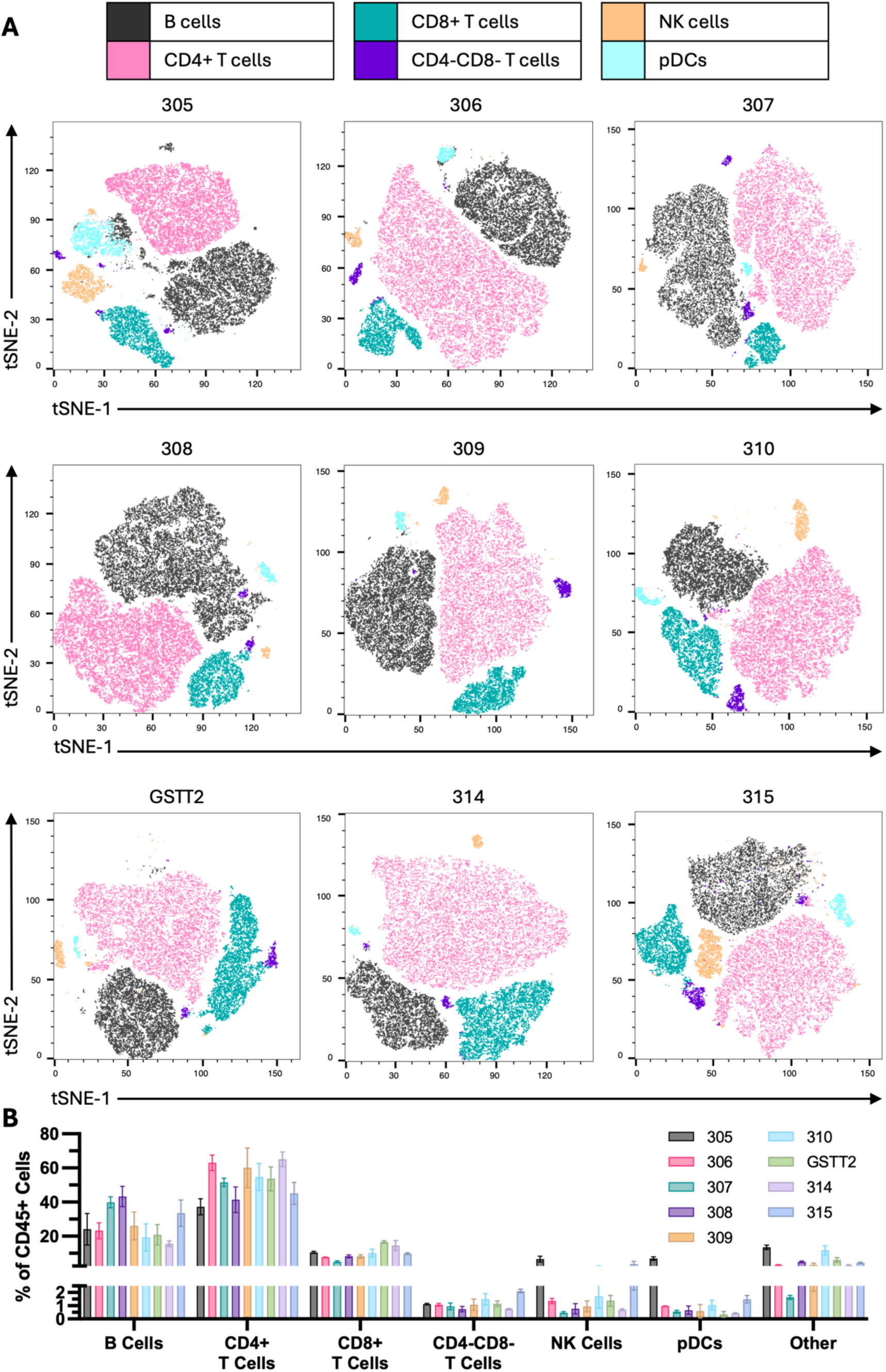
FNA acquires major immune cell populations across all patient-derived reactive ALN. (**A**) Each panel is a representative tSNE from one reactive ALN from each of nine patients, showing the immune cell populations collected in the FNA. (**B**) Graph showing inter-patient variability of the immune cell proportions collected from ALN using FNA.

**Figure S4.**
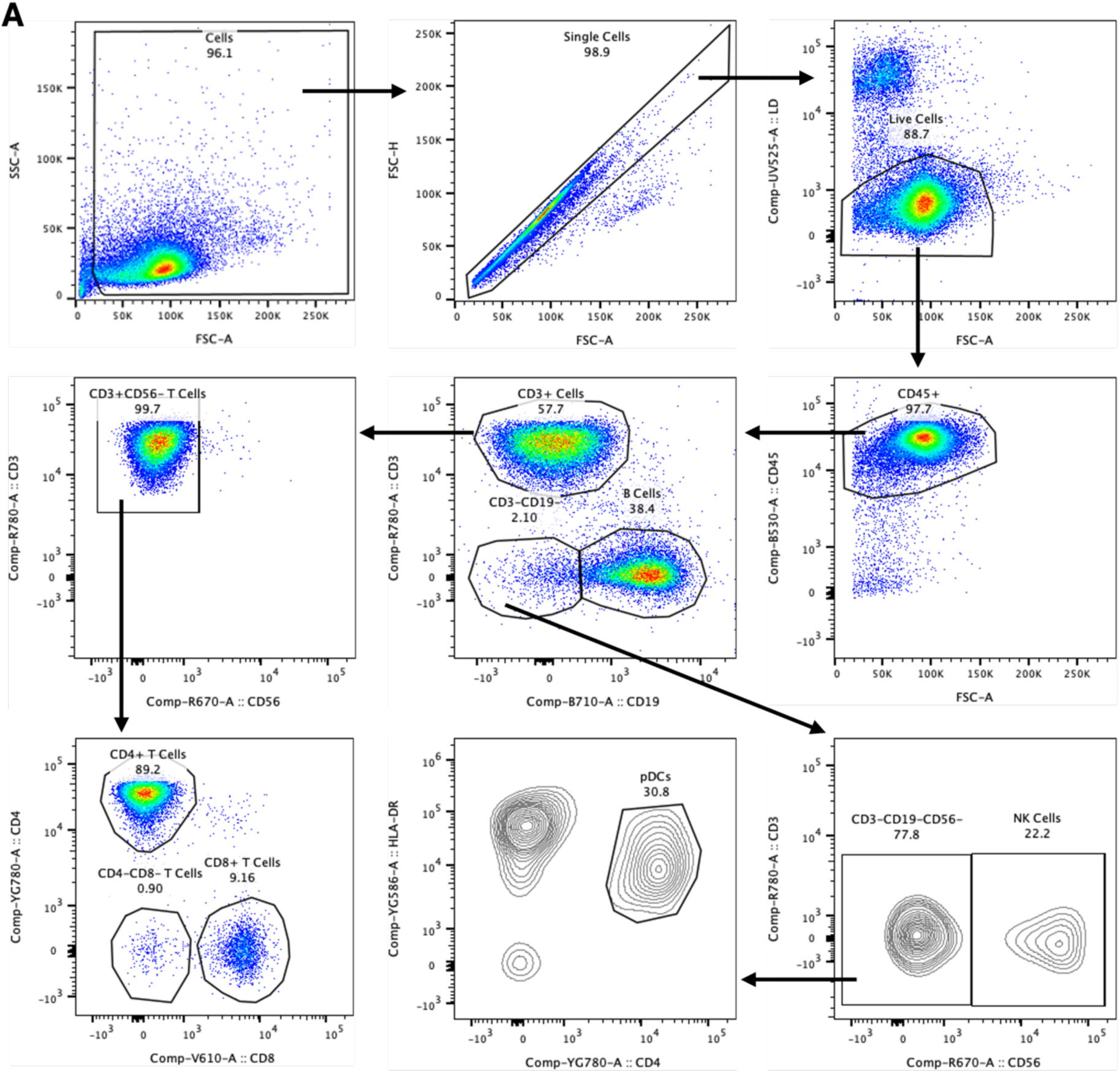
Gating strategy to define major immune cell populations in FNA samples from breast cancer patient-derived reactive ALN.

**Figure S5.**
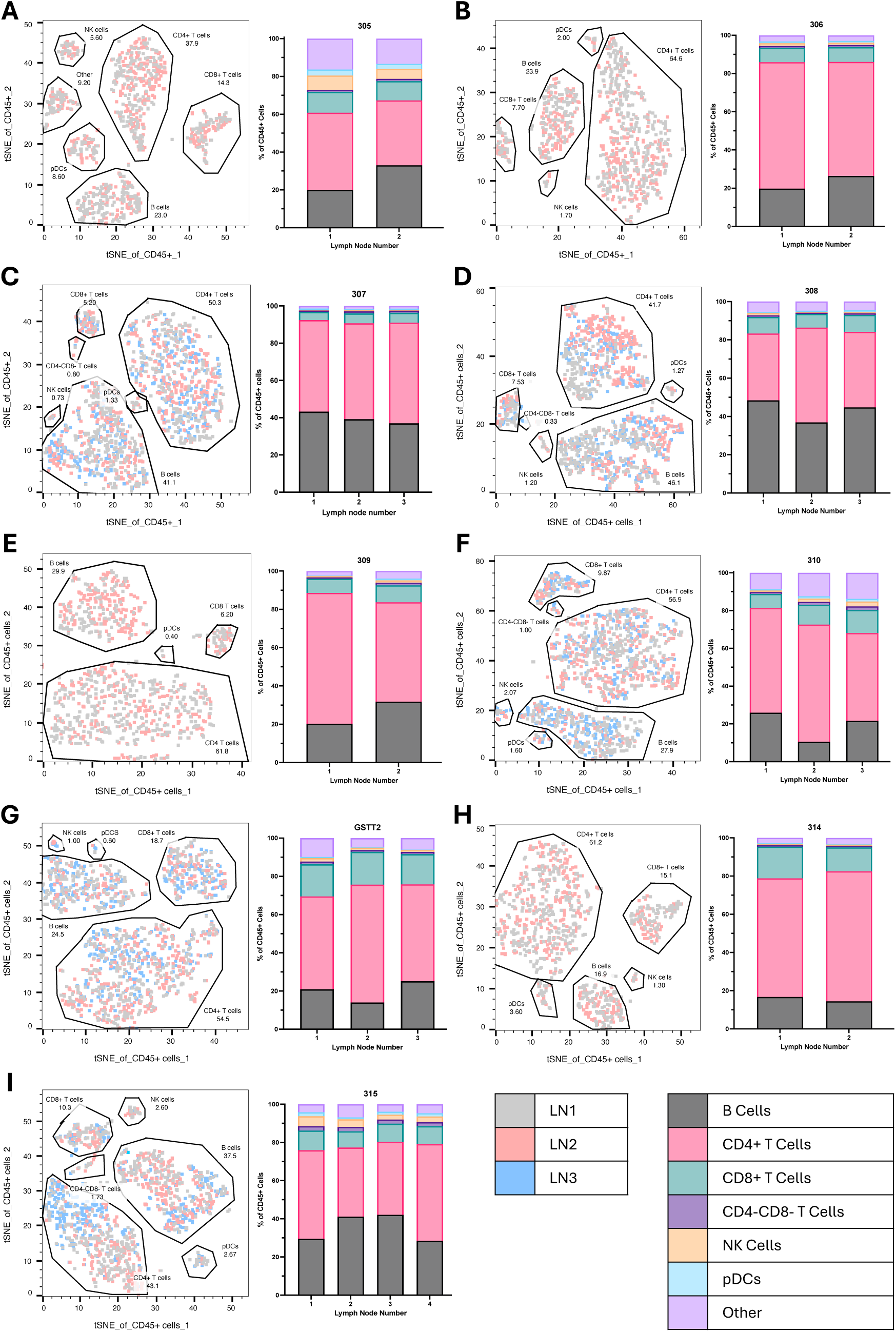
One reactive LN is consistently representative of all reactive LN in a patient’s axilla. (**A – I**) represent each of nine patients that had at least two reactive LNs sampled using FNA in the axilla. TSNEs represent clustering of immune cells from reactive ALN sampled from the same patient. Stacked bar charts show the proportions of each immune cell population from each ALN.

**Figure S6.**
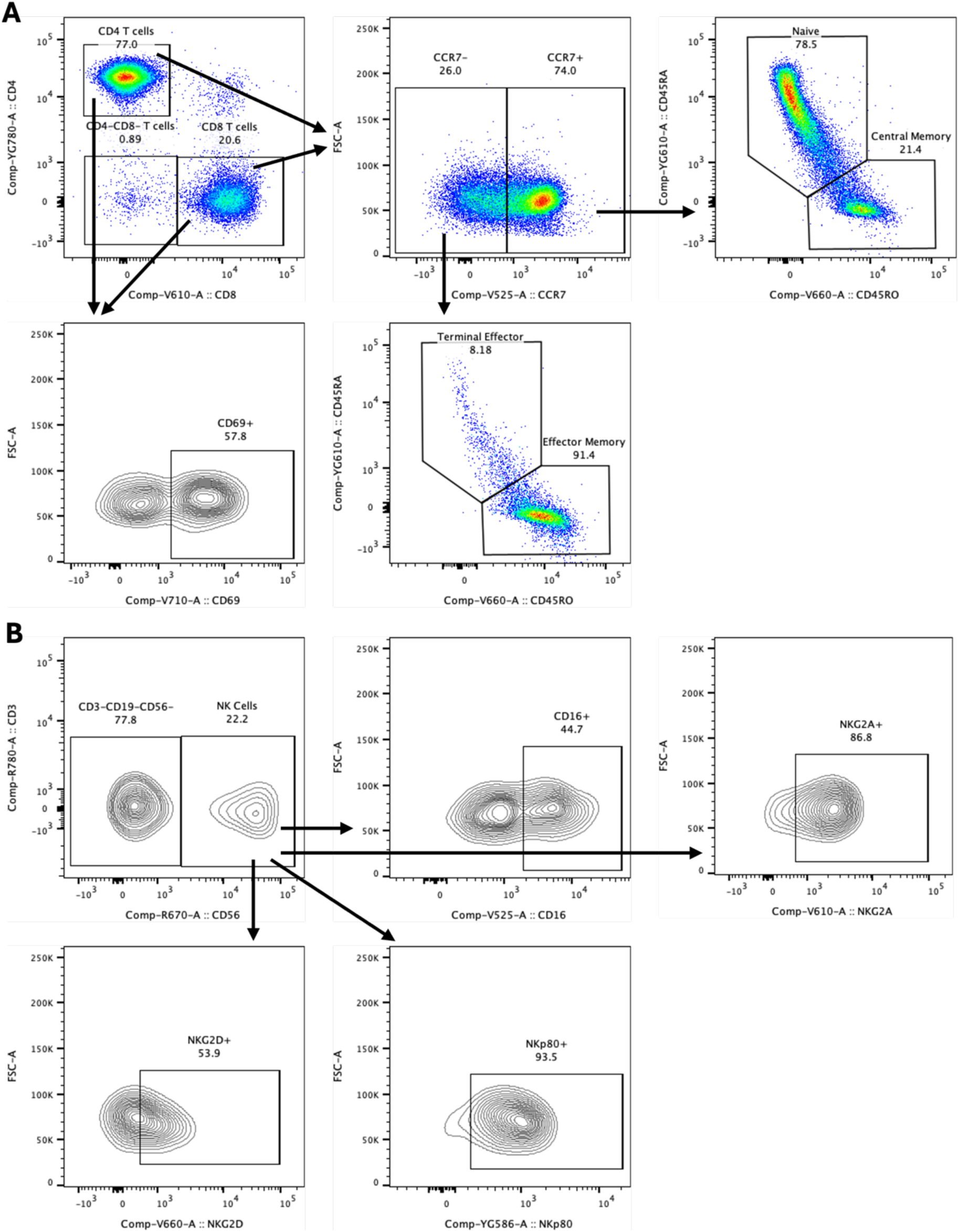
Gating strategies to define immune cell subsets and receptor expression. (**A**) Gating strategy to define CD4+ and CD8+ T cell subsets (naïve, terminal effector, central memory and effector memory), and CD69 and CCR7 expression. (**B**) Gating strategy to determine receptor expression (CD16, NKG2A, NKG2D and NKp80) on NK cells.

**Figure S7.**
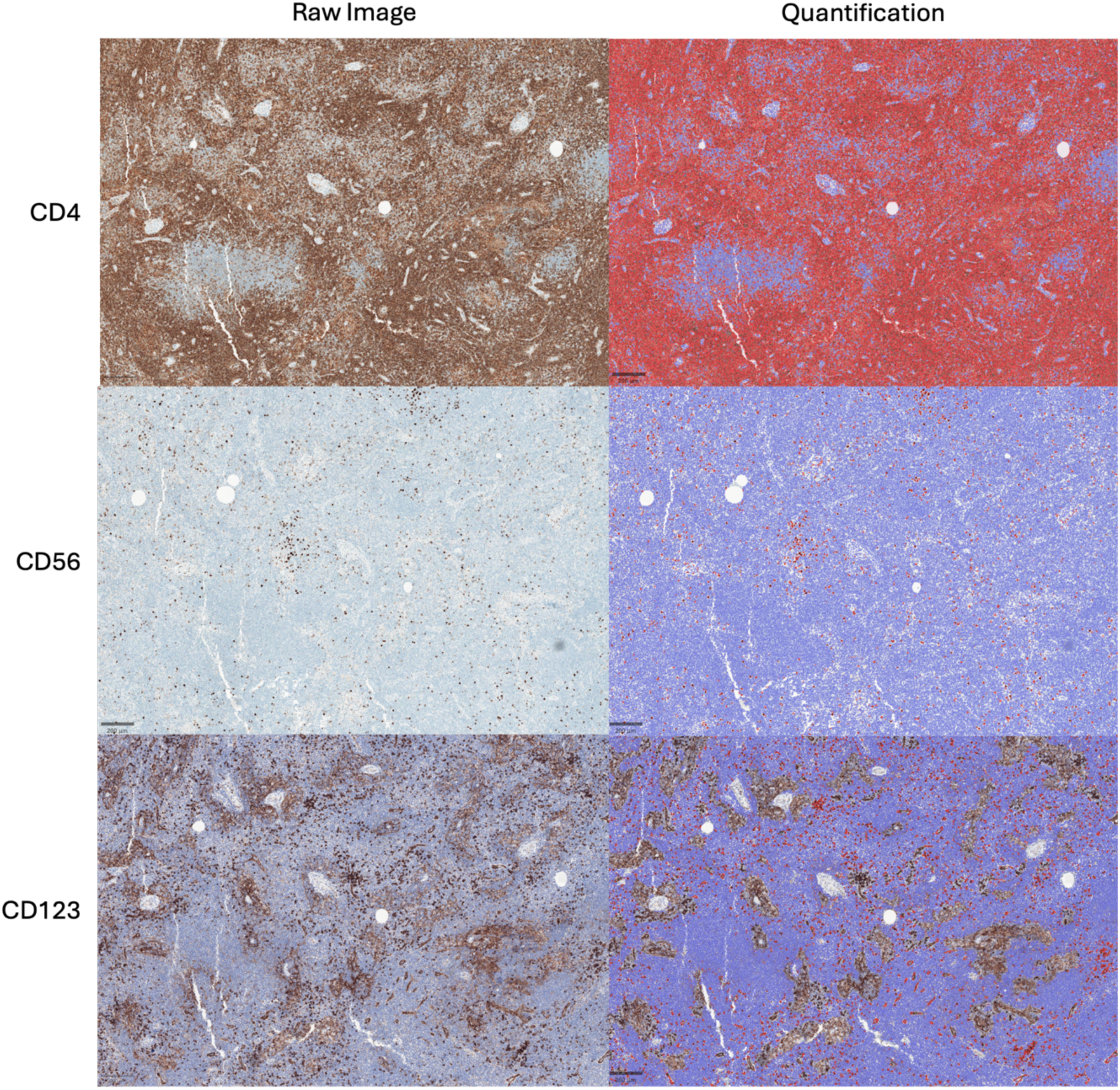
Quantification of CD4^+^, CD56^+^ or CD123^+^ cells in IHC images. Red (positive) and blue (negative). Scale bars 200 µm.

**Table S1.** Anti-mouse antibodies used to compare FNA sampling to whole-lymph node digestion.

| Target | Clone | Fluorochrome | Supplier |
| --- | --- | --- | --- |
| Zombie UV Fixable Viability Dye | N/A | Zombie UV | Biolegend |
| CD3e | 145-2C11 | BUV395 | BD Horizon™ |
| CD4 | RM4-5 | BV510 | Biolegend |
| CD8a | 53-6.7 | PE/Cyanine5 | Biolegend |
| CD11b | M1/70 | APC/Cyanine7 | Biolegend |
| CD11c | N418 | BV650 | Biolegend |
| CD16/32 | 93 | N/A | Biolegend |
| CD19 | 6D5 | BV785 | Biolegend |
| CD45 | 30-F11 | FITC | Biolegend |
| F4/80 | BM8 | PE/Cyanine7 | Biolegend |
| NK-1.1 | PK136 | PE | Biolegend |

**Table S2.** Anti-human antibodies used to define immune cell populations from patient ALN.

| Target | Clone | Fluorochrome | Supplier |
| --- | --- | --- | --- |
| Zombie UV Fixable Viability Dye | N/A | Zombie UV | Biolegend |
| Human TruStain FcX™ Block | N/A | N/A | Biolegend |
| CD3 | HIT3a | APC/Cyanine7 | Biolegend |
| CD4 | SP35 | None | Roche |
| CD4 | SK3 | PE/Cyanine7 | Biolegend |
| CD8a | RPA-T8 | BV605 | Biolegend |
| CD16 | 3G8 | BV510 | Biolegend |
| CD19 | SJ25C1 | PerCP/Cyanine5.5 | Biolegend |
| CD19 | SJ25C1 | APC/Cyanine7 | Biolegend |
| CD45 | HI30 | FITC | Biolegend |
| CD45RA | HI100 | PE/Dazzle | Biolegend |
| CD45RO | UCHL1 | BV650 | Biolegend |
| CD56 | 123C3 | None | Dako |
| CD56 | 5.1H11 | APC | Biolegend |
| CD69 | FN50 | BV711 | Biolegend |
| CD123 | BR4MS | None | Cell Signaling Technology |
| CD197 (CCR7) | G043H7 | PE/Dazzle | Biolegend |
| CD197 (CCR7) | G043H7 | BV510 | Biolegend |
| HLA-DR | L243 | PE | Biolegend |
| NKG2A | S19004C | BV605 | Biolegend |
| NKG2D | 1D11 | BV650 | Biolegend |
| NKp80 | 5D12 | PE | Biolegend |

## Notes

### Competing Interest Statement

The authors have declared no competing interest.

